# Processing of a single ribonucleotide embedded into DNA by human nucleotide excision repair and DNA polymerase η

**DOI:** 10.1101/699587

**Authors:** Akira Sassa, Haruto Tada, Ayuna Takeishi, Kaho Harada, Megumi Suzuki, Masataka Tsuda, Hiroyuki Sasanuma, Shunichi Takeda, Kaoru Sugasawa, Manabu Yasui, Masamitsu Honma, Kiyoe Ura

## Abstract

DNA polymerases often incorporate non-canonical nucleotide, i.e., ribonucleoside triphosphates into the genomic DNA. Aberrant accumulation of ribonucleotides in the genome causes various cellular abnormalities. Here, we show the possible role of human nucleotide excision repair (NER) and DNA polymerase η (Pol η) in processing of a single ribonucleotide embedded into DNA. We found that the reconstituted NER system can excise the oxidized ribonucleotide on the plasmid DNA. Taken together with the evidence that Pol η accurately bypasses a ribonucleotide, i.e., riboguanosine (rG) or its oxidized derivative (8-oxo-rG) *in vitro*, we further assessed the mutagenic potential of the embedded ribonucleotide in human cells lacking NER or Pol η. A single rG on the *supF* reporter gene predominantly induced large deletion mutations. An embedded 8-oxo-rG caused base substitution mutations at the 3’-neighboring base rather than large deletions in wild-type cells. The disruption of *XPA*, an essential factor for NER, or Pol η leads to the increased mutant frequency of 8-oxo-rG. Furthermore, the frequency of 8-oxo-rG-mediated large deletions was increased by the loss of Pol η, but not XPA. Collectively, our results suggest that base oxidation of the embedded ribonucleotide enables processing of the ribonucleotide via alternative DNA repair and damage tolerance pathways.

## 1. Introduction

DNA replication is essential for maintaining genetic information in living organisms. DNA polymerases (Pols) specifically utilize the DNA precursors deoxynucleoside triphosphates (dNTPs) during replication; however, they often incorporate ribonucleoside triphosphates (rNTPs) into DNA because of the much higher concentration of rNTPs than that of dNTPs in the cellular nucleotide pool^1,2^. It has been suggested that millions of ribonucleotides are incorporated into the genome of eukaryotic cells^3,4^. The aberrant accumulation of ribonucleotides in the genome leads to Aicardi–Goutières syndrome (AGS), the severe autoimmune disease, and tumorigenesis^5–7^.

In eukaryotes, ribonucleotides embedded into DNA are primarily repaired by RNase H2-initiated ribonucleotide excision repair (RER)^8^. RNase H2 cleaves the 5’-side of the ribose sugar backbone, followed by excision by a flap endonuclease, strand displacement synthesis by Pol δ or ε, and nick sealing catalyzed by DNA ligase I. The inactivation of the canonical RER leads to various cellular abnormalities including the chronic activation of DNA damage response, aberrant innate immune activation, and epigenetic perturbations^3,7,9–12^. In the absence of RNase H2, the ribonucleotides are repaired by an alternative pathway involving DNA topoisomerase 1 (Top1). This Top1-mediated repair pathway is demonstrated to be associated with mutagenic consequences^13–15^. In yeast cells lacking RNase H2, an embedded ribonucleotide induces 2-to 5-bp deletion mutations at short tandem repeat sequences via Top1-mediated ribonucleotide processing^14^. However, whole-exome sequencing detected no such mutations in the genome of tumor cells derived from RNase H2 deficient mice^5^. The mutagenic potential of ribonucleotides embedded into DNA in mammalian cells remains to be elucidated.

Cellular DNA/RNA and its precursors are subject to oxidation from endogenous and exogenous sources of reactive oxygen species. The exposure of DNA to oxidative stress has been shown to promote the formation of a substantial amount of ribonucleotides and oxidized base damages *in vitro* and *in vivo*^16^. 7,8-Dihydro-8-oxo-riboguanosine-5’-triphosphate (8-oxo-rGTP), the oxidized form of rNTPs, can be generated in the nucleotide pool by the action of oxygen radicals^17^. 8-Oxo-rGTP is also utilized as the substrate during DNA synthesis by Pols^18–20^. Thus, the modified ribonucleotides could exist in DNA via the oxidation of the embedded ribonucleotide and/or the insertion of damaged rNTPs during DNA replication. Base oxidation in embedded ribonucleotide inhibits RNase H2-mediated excision repair^21,22^. Furthermore, the base excision repair (BER) pathway, involved in the repair of 7,8-dihydro-8-oxo-deoxyguanosine (8-oxo-dG), cannot excise the oxidized ribonucleotide 7,8-dihydro-8-oxo-riboguanosine (8-oxo-rG) in the DNA^21,22^ Therefore, a modified ribonucleotide would exhibit greater mutagenic potential compared with unmodified ribonucleotide in cells. Otherwise, an alternative repair pathway may participate in the repair of a damaged ribonucleotide. A possible mechanism is nucleotide excision repair (NER) involved in the removal of various helix-distorting DNA lesions from DNA^23^. A previous study has revealed that NER proteins derived from thermophilic eubacteria can excise ribonucleotides embedded into DNA^24^. However, another study has suggested that a ribonucleotide embedded into DNA is a poor substrate for NER function *in vitro* in both human and *Escherichia coli (E. coli)*^25^. Based on these findings, the involvement of NER in ribonucleotide removal in cells remains controversial.

Ribonucleotides that escape from repair can interfere with DNA replication. *In vitro* studies have revealed that single and multiple ribonucleotides in the template DNA influence DNA synthesis mediated by Pols in yeast and humans^2,26^. Furthermore, the presence of 8-oxo-rG in DNA strongly hinders the primer extension reaction catalyzed by human replicative Pol α^21^. To counteract these replication-blocking lesions, organisms possess specialized Pols that carry out translesion DNA synthesis (TLS) past DNA lesions. Pol η is one of TLS Pols that can bypass various DNA damages, including ultraviolet (UV) light-induced cyclobutane pyrimidine dimers^27^. Our *in vitro* experiments revealed that Pol η efficiently and accurately bypassed undamaged and damaged ribonucleotides (rG and 8-oxo-rG respectively) in a more error-free manner compared with 8-oxo-dG^21^. Thus, it is of interest to investigate whether TLS Pols exert protective effects against the mutagenicity of incorporated ribonucleotides in cells.

In the current study, we investigated the possibility whether the reconstituted NER system can excise the ribonucleotide from the DNA. On the basis of *in vitro* analyses, we also examined the mutagenic potential of an embedded ribonucleotide in human lymphoblastoid cells lacking xeroderma pigmentosum group A (XPA), an essential factor for NER, and Pol η using a *supF* shuttle vector containing a single dG rG, 8-oxo-rG, or 8-oxo-dG Based on these biochemical and cell culture investigations, our findings provide new mechanistic insights underlying ribonucleotide-induced mutagenesis and the protective roles of DNA repair and tolerance processes in mammals.

## Results

### Reconstituted NER can excise an embedded oxidized ribonucleotide

To test the possibility that NER is involved in the ribonucleotide removal from DNA, we analyzed whether the human cell-free NER system reconstituted with purified protein factors possesses the ribonucleotide excision activity against a single rG or 8-oxo-rG in the plasmid DNA (Fig. 1a). As shown in Figure 1b, single rG was a poor substrate for human NER, a finding consistent with the previous report^25^. Conversely, 8-oxo-rG was recognized and excised by NER machinery. In addition, the incision activity of NER against 8-oxo-rG was greater than that against 8-oxo-dG (Fig. 1c). The ladder pattern of the excised products was also slightly different between 8-oxo-dG and 8-oxo-rG (Fig. 1b), which can be due to the difference in the sugar pucker conformation of deoxyribose and ribose that may affect the recognition and excision processes of NER^28^. Because the efficiency of NER in excising the ribonucleotide was lower compared with the canonical substrate for NER, i.e., UV-induced 6-4 photoproducts (Fig. 1b) (Supplementary Fig. S1), we further assessed the cellular impact of the NER deficiency on the mutagenic potential of an embedded ribonucleotide in the following section.

**Fig. 1.**
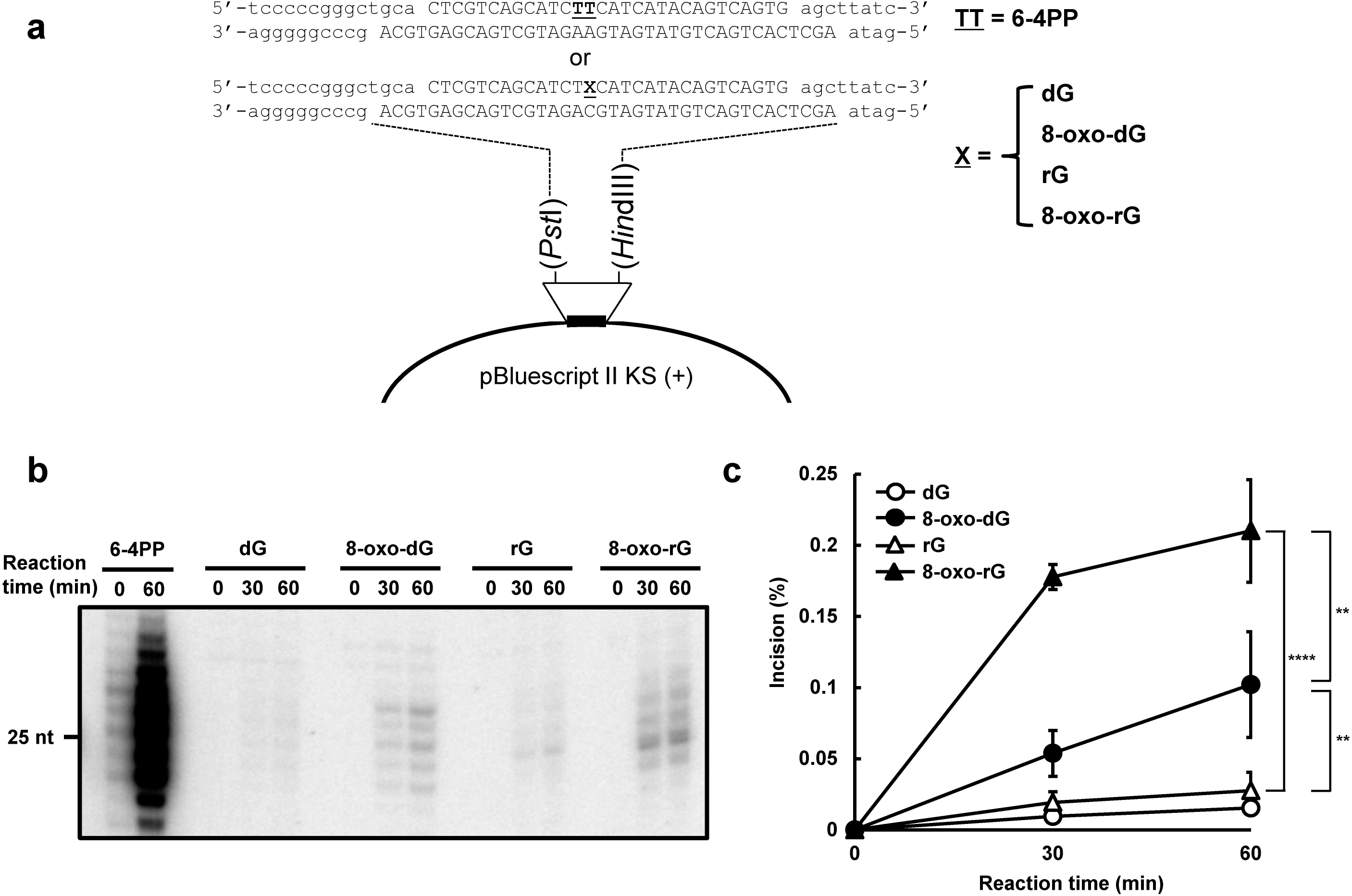
The dual incision activity of human nucleotide excision repair on duplex plasmid DNA containing a single ribonucleotide. (a) The DNA substrate used for the *in vitro* dual incision assay. The substrate plasmid harbored a single 6-4PP or dG 8-oxo-dG, rG, or 8-oxo-rG at a defined position in the DNA sequence between PstI and *HindIII* sites of pBluescript II KS(+) as previously described^53^. (b) *In vitro* dual incision assays were performed using purified NER proteins and internally ^32^P-labeled DNA substrates containing a 6-4PP, dG, 8-oxo-dG, rG, or 8-oxo-rG as described in the Methods. DNA samples were subjected to 10% denaturing polyacrylamide gel electrophoresis followed by autoradiography. The full-length gel is shown in Supplementary Figure S1. (c) Quantification of the products formed by NER-mediated dual incision reaction. Values are presented as mean ± S.E. of three independent experiments. Significant differences are indicated by asterisks (**P < 0.01, ****P < 0.0001 by two-way ANOVA).

### Experimental design to determine the mutagenic potential of an embedded ribonucleotide in human cells

To examine the mutagenic potential of an embedded ribonucleotide in human cells, double-stranded plasmids containing rG or 8-oxo-rG were constructed as the model substrates. Ribonucleotides were inserted at position 112 (G^112^) on *supF* (Fig. 2a). The cThe stable expression oontrol plasmids harboring dG or 8-oxo-dG at the same position were also synthesized to clarify the effect of the ribose sugar backbone on mutations. The phenotype of cells used in this study was confirmed prior to the experiments; *XPA^−/−^* and *POLH^−/−^* cells were hypersensitive to UV exposure (Fig. 2b), as previously described^29,30^. The stable expression of *XPA* and *POLH* in each knock-out cells, i.e., “*XPA^−/−^* + *XPA*” and “*POLH^−/−^ + POLH*”, rescued the phenotype, which was then utilized for the complementation experiment as described in the following section.

**Fig. 2.**
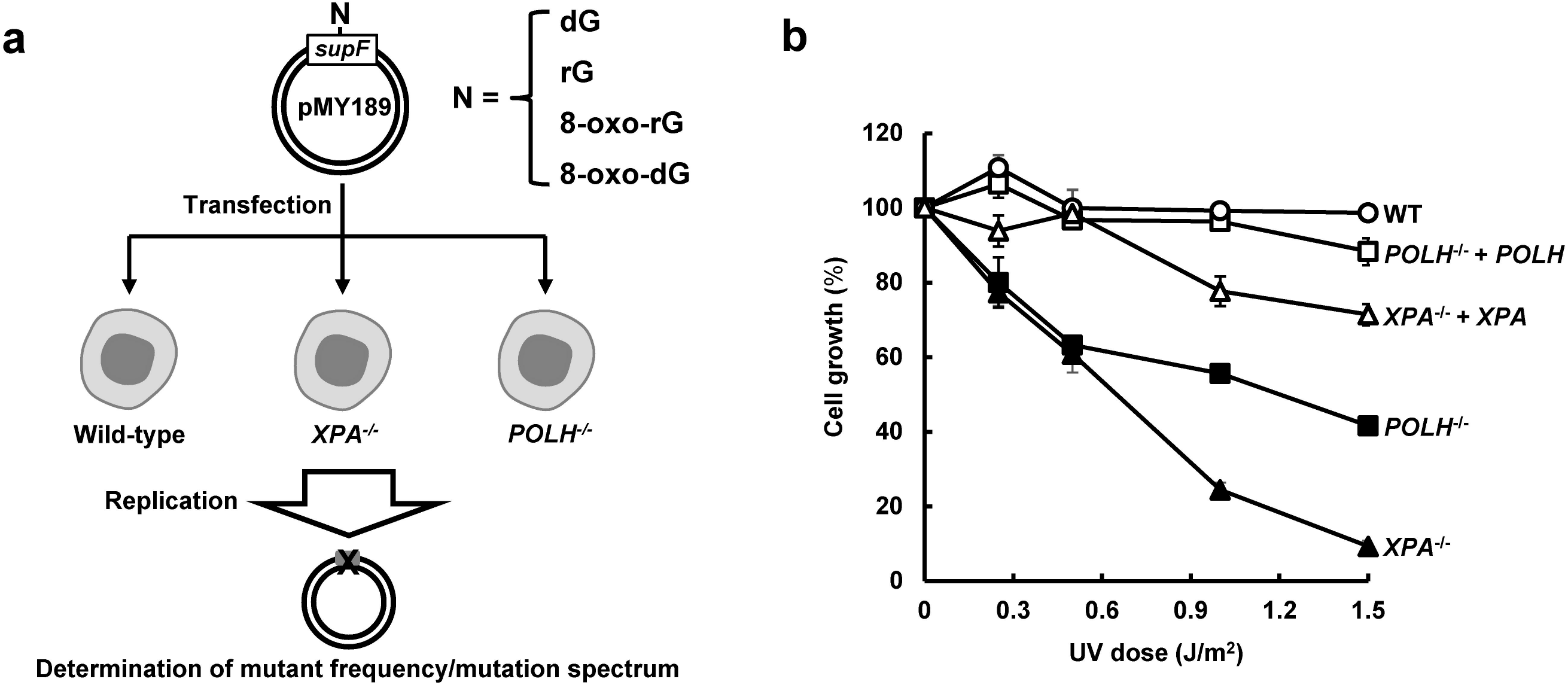
Analysis of ribonucleotide-induced mutagenesis in human lymphoblastoid cells. (a) Schematic diagram of the determination of mutant frequency and mutation spectrum using the *supF* shuttle vector. The closed circular double-stranded DNA, containing a single dG, rG, 8-oxo-rG, or 8-oxo-dG in *supF*, was transfected into WT, *XPA^−/−^*, and *POLH^−/−^* cells. After incubation for 48 h, the propagated plasmids were extracted from cells and introduced into KS40/pOF105 indicator strain. The mutant frequency and mutation spectrum were determined as described in the Methods. (b) Cell growth of WT (open circles), *XPA^−/−^* (closed triangles), *POLH^−/−^* (closed squares), *XPA^−/−^* + *XPA* (open triangles), and *POLH^−/−^* + *POLH* (open squares) cells after exposure to UVC light. Values are presented as mean ± S.E. of at least two independent experiments.

### A single ribonucleotide predominantly induces large deletions in human cells

First, the plasmid DNA containing a single rG was transfected and replicated in wild-type (WT), *XPA^−/−^*, and *POLH^−/−^* cells. Plasmids replicated in cells were then recovered and introduced into KS40/pOF105 indicator strain, which was plated on titer and selection plates to calculate the frequencies of *supF* mutants. The mutant frequency induced by rG was comparable between the cells (3.9 ± 0.19, 3.6 ± 0.76, and 3.7 ± 0.056 × 10^−3^ in WT, *XPA^−/−^*, and *POLH^−/−^* cells, respectively), which was approximately four-fold higher than the control dG (0.88 ± 0.35, 0.79 ± 0.30, and 0.63 ± 0.12 × 10^−3^ in WT, *XPA^−/−^*, and *POLH^−/−^* cells, respectively) (Fig. 3a and b). We further analyzed the mutation spectrum induced by rG. Interestingly, large deletions were the predominant mutation (67%, 75%, and 82% in WT, *XPA^−/−^*, and *POLH^−/−^* cells, respectively) (Fig. 4b) (Supplementary Table S1). The length of deletions varied from 10 to 244 bp, with the majority of them being >100 bp in all cell lines (Supplementary Table S2). Notably, no sequence homology was found at each deletion junction. Because the frequency of large deletions observed was only approximately 10% in the control experiments involving dG (Fig. 4a) (Supplementary Table S1), large deletion mutations were specifically induced by rG embedded into the DNA.

**Fig. 3.**
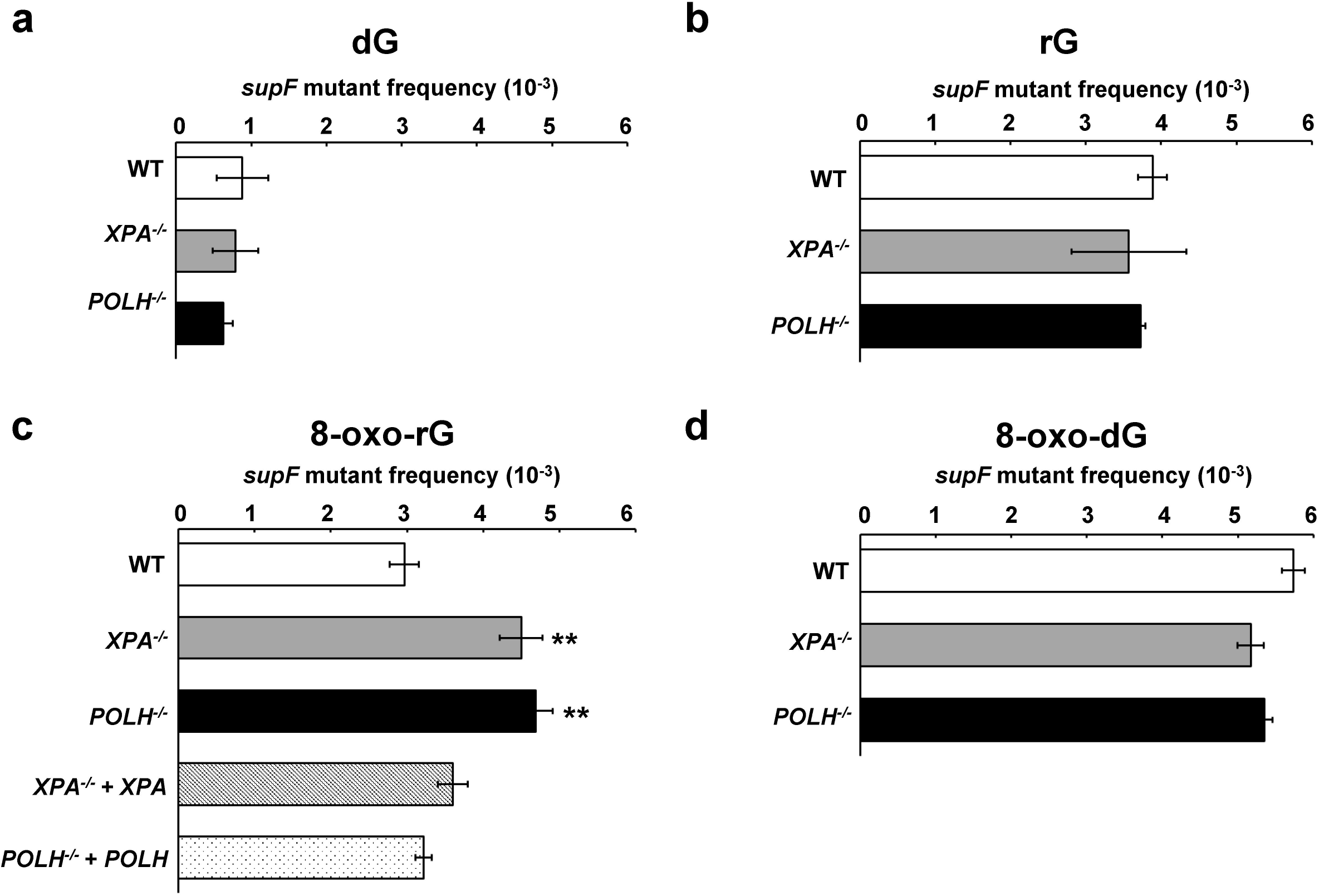
Frequencies of *supF* mutants in a shuttle vector containing (a) dG, (b) rG, (c) 8-oxo-rG, and (d) 8-oxo-dG propagated in human cells. Data are expressed as mean ± S.E. of 2-5 independent experiments. ** Significant difference between the assessed cells and WT cells; P < 0.01 (Dunnett’s Multiple Comparison Test).

**Fig. 4.**
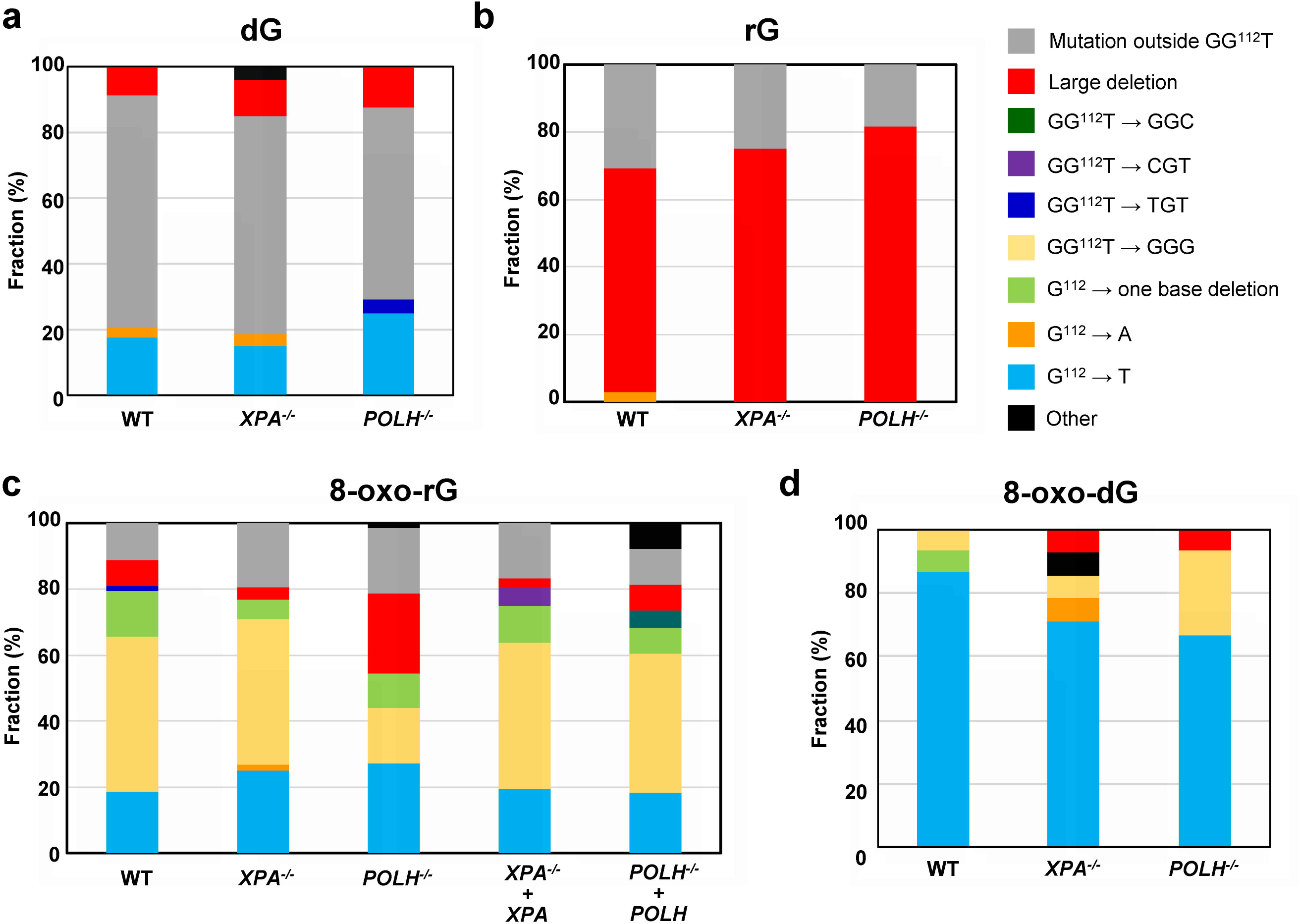
Mutation spectrum of (a) dG, (b) rG, (c) 8-oxo-rG, and (d) 8-oxo-dG in *supF* plasmids propagated in wild-type, *POLH^−/−^*, and *XPA^−/−^* cells. The number of mutations were summarized in Supplementary Table S1.

### Disruption of *XPA* or *POLH* enhances the mutagenic potential of an oxidized ribonucleotide

Next, we assessed whether base oxidation in a ribonucleotide affected its mutagenic potential in cells. The mutant frequency induced by 8-oxo-rG (3.0 ± 0.19 × 10^−3^) was not higher than that induced by rG (3.9 ± 0.19 × 10^−3^) and 8-oxo-dG (5.7 ± 0.15 × 10^−3^) in WT cells (Fig. 3b and c). In contrast with 8-oxo-dG that exclusively induced G:C to T:A transversion mutations at the position G^112^ (87%) (Fig. 4d) (Supplementary Table S1), 8-oxo-rG preferentially induced base-substitution mutations at a 3’-base neighboring the 8-oxo-rG site, i.e., GG^112^T to GGG (46%), followed by G:C to T:A (19%) and one-base deletions (14%) at the site of 8-oxo-rG (Fig. 4c). The frequency of large deletions induced by 8-oxo-rG was 7.9%, which was lower than that induced by rG (67%) (Fig. 4b). As some extent of G:C to T:A mutation (18%) was also induced by the control dG (Fig. 4a), mutations specifically induced by 8-oxo-rG were GG^112^T to GGG and one-base deletions, rather than large deletions and G:C to T:A transversion mutations. GG^112^T to GGG mutation was also observed with 8-oxo-dG in WT cells (6.7%) (Fig. 4d), suggesting that this mutation depends on the sequence context around G^112^.

To investigate the role of NER and TLS Pol on the mutagenic events caused by base oxidation in embedded ribonucleotide, we examined the mutant frequency and mutation spectrum of 8-oxo-rG in *XPA^−/−^* and *POLH^−/−^* cells. The mutant frequency was significantly increased in *XPA^−/−^* (4.5 ± 0.28 × 10^−3^) and *POLH^−/−^* (4.7 ± 0.22 × 10^−3^) compared with that in WT cells (3.0 ± 0.19 × 10^−3^) (Fig. 3c). The loss of *XPA* did not alter the mutation spectrum induced by 8-oxo-rG compared with that in WT cells (Fig. 4c) (Supplementary Table S1). In contrast, the loss of *POLH* resulted in an increased number of 8-oxo-rG-mediated large deletions (24%) in comparison with WT cells (7.9%). These deletions were mainly observed at a position 5 □-upstream from the site of 8-oxo-rG (Supplementary Table S2). Accordingly, the number of GG^112^T to GGG mutation was decreased in *POLH^−/−^* (17%) as compared with WT cells (46%). To verify whether the increased mutagenic potential of 8-oxo-rG was due to the deficiency of *XPA* or *POLH*, we introduced the plasmid containing 8-oxo-rG into *XPA^−/−^* + *XPA* and *POLH^−/−^* + *POLH* cells. The frequency of the 8-oxo-rG-induced *supF* mutant was 3.6 ± 0.11 × 10^−3^ and 3.2 ± 0.11 × 10^−3^ in *XPA^−/−^* + *XPA* and *POLH^−/−^* + *POLH* cells, respectively, which was comparable with that in WT cells (3.0 ± 0.19 × 10^−3^) (Fig. 3c). The expression of *POLH* in *POLH^−/−^* cells also decreased the frequency of large deletions (7.9%) (Fig. 4c) (Supplementary Table S1 and S2). It is noted that the disruption of *XPA* or *POLH* did not alter the mutant frequency of 8-oxo-dG (5.7 ± 0.15 × 10^−3^, 5.2 ± 0.17 × 10^−3^ and 5.4 ± 0.10 × 10^−3^ in WT, *XPA^−/−^* and *POLH^−/−^* cells, respectively) (Fig. 3d). The predominant mutation induced by 8-oxo-dG was G:C to T:A transversion mutations in all cells (87%, 71%, and 67% in WT, *XPA^−/−^*, and *POLH^−/−^* cells, respectively) (Fig. 4d) (Supplementary Table S1). The overall mutation spectrum was slightly altered in *POLH^−/−^* cells; the number of GG^112^T to GGG was increased (Table S1), which might be because of the TLS event across 8-oxo-dG by other Pols in the absence of Pol η.

## Discussion

The absence of RER leads to the accumulation of ribonucleotides in the genome, which is associated with increased genomic instability^31^. Although several studies have reported the importance of RER for maintaining genome integrity in higher eukaryotes^3,32^, the mutational specificity of embedded ribonucleotides remains largely unclear in mammals. To visualize mutations induced by a ribonucleotide that is readily repaired by RER in the genome, we employed the shuttle vector system that can rapidly propagate in cells. Our results suggest that an embedded rG can cause the deleterious mutations in human cells. On the other hand, its oxidative derivative 8-oxo-rG did not display the enhanced mutagenic potential compared with rG and 8-oxo-dG, which can be explained by the alternative repair and damage tolerance pathways based on *in vitro* investigations using the reconstituted NER and Pol η.

It has been controversial whether NER can recognize a small lesion such as 8-oxo-dG-and ribonucleotide-containing DNA as the substrate^24,25,30,33–35^. The *in vitro* NER reconstitution experiments in this study implicated that 8-oxo-rG is the preferable substrate for human NER compared with rG and 8-oxo-dG in the plasmid DNA (Fig. 1c). Further, our results revealed that the loss of the NER factor *XPA* does not alter the mutant frequency and mutation spectra induced by rG and 8-oxo-dG, but significantly enhanced the mutagenic potential of 8-oxo-rG (Fig. 3). Hence, these observations suggest that the oxidation of guanine base in ribonucleotide enables ribonucleotide removal by human NER.

In a previous *in vitro* study, both replicative and TLS Pols bypassed rG in an error-free manner^21^. Conversely, the oxidation of a ribonucleotide, i.e., 8-oxo-rG, on the template DNA strongly inhibited DNA synthesis by Pols and contributed to miscoding events. Because 8-oxo-rG also could not be repaired by either RER or BER^21,22^, we hypothesized that such modification of the ribonucleotide enhances the mutagenic potential in cells. Contrary to the initial hypothesis, it seems that 8-oxo-rG did not display greater mutagenic potential than rG in WT cells (Figure 3b and c). The frequency of large deletions induced by rG (67%) was much higher than that induced by 8-oxo-rG (7.9%) (Fig. 4b and c). These observations may be attributed to the recognition of the damaged ribonucleotide by the alternate repair and/or damage tolerance pathways. While 8-oxo-rG is the replication blocking lesion, Pol η rapidly and accurately bypasses the lesion^21^. Along with the *in vitro* study, our results suggest that the oxidized ribonucleotide on the template DNA is recognized by the DNA damage tolerance pathway and Pol η performs TLS in cells. Interestingly, the proportion of GG^112^T to GGG mutation caused by 8-oxo-rG was decreased in *POLH^−/−^* cells (Fig. 4c), indicating that such mutation is induced during Pol η-mediated TLS across 8-oxo-rG. Although the structural impact of the presence of 8-oxo-rG in DNA is still unclear, the mutation could be because of the sequence specific misalignment induced by 8-oxo-rG For example, the nucleotide incorporated opposite T^113^ is encoded by 5’-neighboring 8-oxo-rG through the misalignment-mediated nucleotide insertion followed by realignment of the template DNA during TLS. It has been suggested that such mutations due to misalignment can be induced by DNA damage at the specific sequence context^36^. While the loss of *POLH* resulted in a three-fold increase in the frequency of large deletion mutations induced by 8-oxo-rG (Fig. 4c), the loss of *XPA* did not enhance the frequency of such deletions despite the increase in the frequency of mutations (Fig. 3c and 4c). Based on these observations, ribonucleotide-induced large deletions may occur in conjunction with DNA synthesis. Our results also imply that an embedded ribonucleotide causes mutations not only in tandem repeat sequences but also in other genomic loci.

In contrast with findings of the current study, previous studies have reported that such large deletion mutations at the non-repetitive sequences were rare in the genome of the yeast strain lacking RER^13–15^. This difference may be because of a distinct repair mechanism activated during the mutagenic processing of the ribonucleotide. During Top1-mediated ribonucleotide processing, Top1 incises the 3’-side of the ribonucleotide and forms a Top1–DNA cleavage complex (Top1cc) between the catalytic tyrosine residue and the 3’-phosphate of the ribonucleotide^14,37^. Top1cc is further reversed by the nucleophilic attack of 2’-OH of the ribonucleotide to generate a 2’,3’-cyclic phosphate and to release Top1 from the complex. The subsequent processing of the released DNA by Top1 can lead to DNA double strand breaks (DSBs)^38^. Possible repair pathways following DSBs formation include homologous recombination (HR) and non-homologous end-joining (NHEJ). The predominant DSB repair pathway is NHEJ throughout the cell cycle in mammalian cells^39^, whereas HR is prevalent especially during the G2/S phase in yeast^40^. This pathway choice may explain the differential mutagenic property of embedded ribonucleotides between yeast and mammalian cells. Interestingly, large deletions induced by rG and 8-oxo-rG were often extended from the site of the ribonucleotide (G^112^) to a position 5 □-upstream in *supF* (Supplementary Table S2). These mutations could result from the Top1-mediated processing of a ribonucleotide in the following manner. The incision of a ribonucleotide by Top1 generates Top1cc. Upon the generation of a 2’,3’-cyclic phosphate-terminated nick and the release of Top1, one possible scenario is the formation of a second Top1cc 5’-upstream of the nick, leaving a gap between 3’-Top1cc and 5’-OH^41,42^. Because Top1 is coupled with DNA replication^43^, the generation of such a gap or nick during replication can lead to DSBs, in which the 3’-end of the strand contains Top1cc. In the present study, the deletion mutations observed toward the 5’ side of *supF* may have occurred via the processing of such complex DSBs, e.g., the resection of the dirty terminus by NHEJ. Reportedly, even a single DNA damage, e.g., cyclobutane pyrimidine dimer, abasic site, and DSB is sufficient to induce large deletion mutations in the episomal and genomic DNA in mammalian cells^44–46^. Our results provide a further insight into the mechanism of ribonucleotide-induced genomic instability and its suppression by alternative factors involving DNA repair and damage tolerance.

Figure 5 shows the probable model of ribonucleotide-induced mutagenesis and the mechanism for resolving a damaged ribonucleotide in DNA. In summary, unrepaired ribonucleotides in DNA can cause severe genomic instability via the formation of deleterious mutations in mammals. Our findings also suggest that NER and Pol η are employed the back-up pathways for suppressing ribonucleotide-induced mutagenesis under circumstances where embedded ribonucleotides cannot be properly resolved. These findings can deepen our understanding of the mechanism underlying genome instability caused by the genomic ribonucleotide accumulation.

**Fig. 5.**
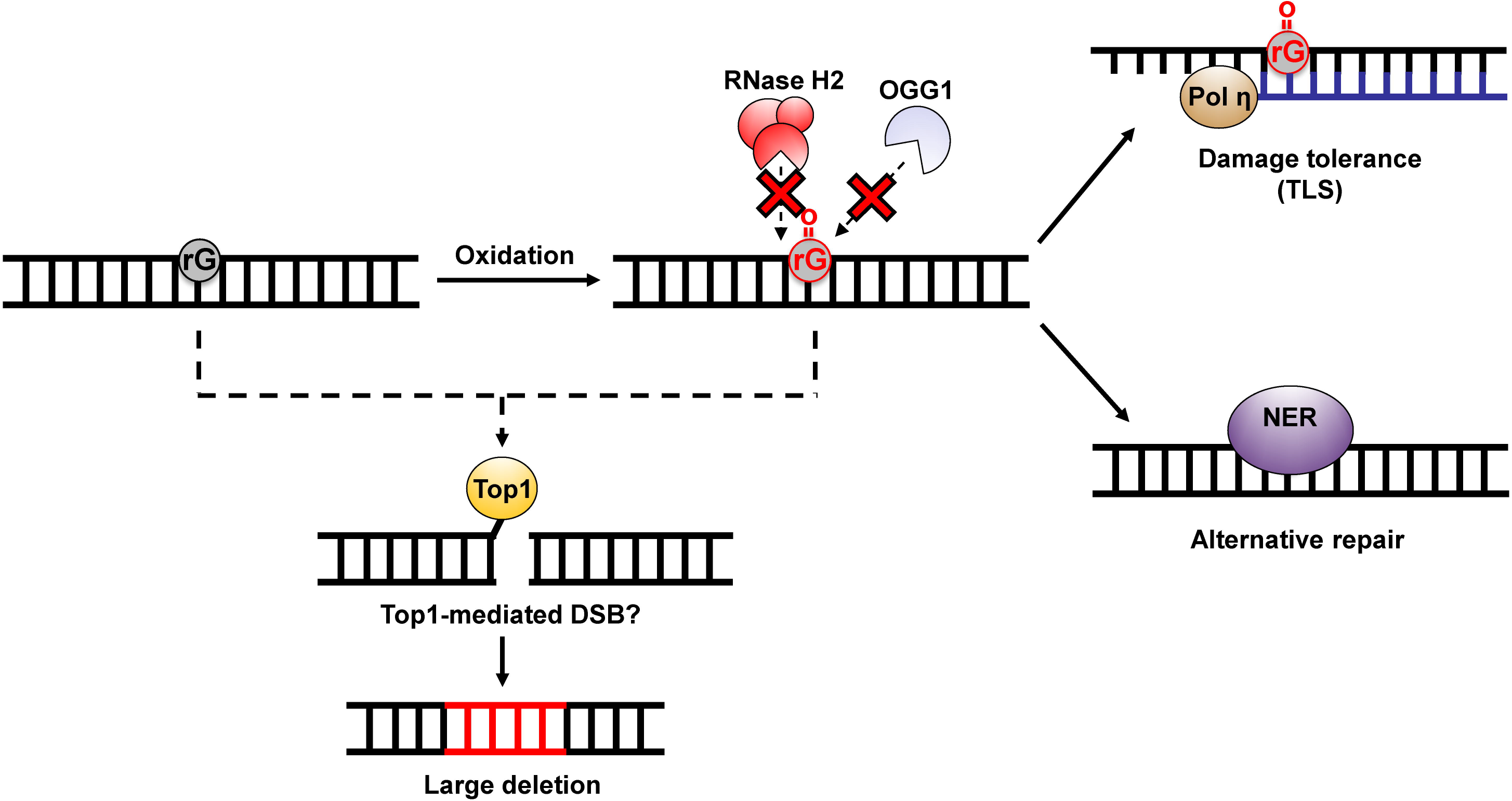
A possible model of the processing of an embedded ribonucleotide and ribonucleotide-induced mutagenesis. 8-Oxo-rG could be generated by the incorporation of oxidized ribonucleoside triphosphates into DNA or by the oxidation of the embedded ribonucleotides. RNase H2 and BER cannot remove 8-oxo-rG^21,22^. NER is involved in the excision of such a damaged ribonucleotide. Alternatively, Pol η can bypass 8-oxo-rG to suppress the mutagenic events induced by the embedded ribonucleotide. The unrepaired ribonucleotide is subject to Top1-mediated processing, leading to the deleterious mutations.

## Methods

### Cell culture

Human lymphoblastoid TSCER2 cells, derived from TK6 cell line^47^, were cultured in RPMI-1640 medium (Nacalai Tesque) supplemented with 200 μg/mL sodium pyruvate, 100 U/mL penicillin, and 100 μg/mL streptomycin, and 10% (v/v) heat-inactivated fetal bovine serum at 37°C in an atmosphere of 5% CO_2_ and 100% humidity. *XPA^−/−^* and *POLH^−/−^* cells were established as described previously^29^.

### Protein preparation

Expression and purification of recombinant NER proteins, Flag-XPC/RAD23B-His^48^, CETN2^49^, TFIIH Core7^50^, Flag-XPA, ERCC1-His/XPF, and RPA^51^, were achieved as previously described. For the preparation of XPG-Flag, High-Five cells (grown in twenty 150-mm culture dishes) were infected with a recombinant baculovirus. After incubation at 27°C for 72 h, the infected cells were collected and washed with ice-cold phosphate-buffered saline. The cell pellet was resuspended in an 8-fold volume of extraction buffer [25 mM Tris-HCl (pH 7.5), 5 mM NaCl, 0.1 mM EDTA, 1.5 mM MgCl_2_, 0.1% Nonidet P-40, 20% glycerol, 1 mM dithiothreitol (DTT), 0.25 mM PMSF, protease inhibitor cocktail (Complete: Roche)]. After incubation on ice for 30 min, the cells were disrupted using Dounce homogenization (20 strokes of pestle A). The cell lysate was then adjusted at 0.3 M NaCl and further incubated on ice for 60 min. A clear extract obtained after centrifugation at 35,000 rpm for 30 min (Beckman 50.2Ti rotor) was loaded onto a HiPrep 16/10 Heparin FF column (GE Healthcare) equilibrated with buffer PET [20 mM sodium phosphate (pH 7.8), 1 mM EDTA, 10% glycerol, 0.01% Triton X-100, 1 mM DTT, 0.25 mM PMSF] containing 0.1 M NaCl. After extensively washing with the same buffer, bound proteins were eluted with buffer PET containing 1 M NaCl. The eluate was then loaded onto an anti-Flag M2 agarose column (1 cm diameter × 5 cm long), which had been equilibrated with buffer PT [20 mM sodium phosphate (pH 7.8), 0.15 M NaCl, 10% glycerol, 0.01% Triton X-100, 1 mM 2-mercaptoethanol, 0.25 mM PMSF). After an extensive wash with the same buffer, the column was inverted and bound proteins were eluted with 10 ml of buffer PT containing 0.1 mg/ml Flag peptide (Sigma). The eluate fractions were dialyzed against buffer A [25 mM Hepes-NaOH (pH 7.5), 0.1 M NaCl, 10% glycerol, 1 mM 2-mercaptoethanol, 0.25 mM PMSF] and loaded onto a HiTrap Q HP column (5 ml, GE healthcare), which was previously equilibrated with the same buffer. Bound proteins were then eluted using a linear NaCl gradient (0.1 to 1 M) in buffer A. The fractions containing XPG-Flag were dialyzed against buffer PET containing 0.1 M NaCl, and loaded onto a Mono S PC 1.6/5 column (GE Healthcare) equilibrated with the same buffer. XPG-Flag was eluted using a linear NaCl gradient (0.1 to 1 M) in buffer PET. The eluted XPG-Flag was dialyzed against buffer PET containing 0.1 M NaCl.

### *In vitro* dual incision assay

The oligonucleotide containing 6-4PP was prepared as described previously^52^. The oligonucleotides containing dG, 8-oxo-dG, rG, or 8-oxo-rG were synthesized and purified by Tsukuba Oligo Service Co., Ltd. (Ibaraki, Japan). The oligonucleotides were purified by polyacrylamide gel electrophoresis. After the purification step, the homogeneity and no detectable degradation of the oligonucleotides were confirmed by the same method prior to use. The internally^32^P-labeled DNA substrates, containing a site-specific lesion, were prepared as previously described^53^. *In vitro* dual incision assays were performed basically according to the previously described procedures^54^. Briefly, the DNA substrates (~1×10^6^ cpm) were incubated at 30°C for the indicated time in 15-μl reactions, each containing 40 mM Hepes-KOH (pH 7.8), 70 mM NaCl, 0.5 mM DTT, 5 mM MgCl_2_, 0.5 mM EDTA, 2 mM ATP, acetylated bovine serum albumin (Ambion: 5.4 μg), Flag-XPA (87.5 ng), ERCC1-His/XPF (15 ng), RPA (100 ng), XPG-Flag (60 ng), TFIIH Core7 (250 ng), Flag-XPC/RAD23B-His (10 ng), and CETN2 (10 ng). The purified DNA samples were subjected to 10% denaturing polyacrylamide gel electrophoresis and were detected using the Typhoon FLA9500 (GE Healthcare) and analyzed with the ImageQuant TL software (GE Healthcare).

### Establishment of *POLH* and *XPA*-complemented cells

The XPA-expressing cell clone (*XPA^−/−^* + *XPA*) was established as previously reported^30^. To construct the *POLH* expression plasmid, the coding region of *POLH* was amplified from the cDNA of TK6 cells by PCR with primers 5’-GAGCTCGGATCCTGAAAAATGGCTACTGGACAGGATC-3’ and 5’-CAGCGGGTTTAAACCTAATGTGTTAATGGCTTAAAAAATGATTCCAATG-3′. The resulting product was digested with *Bam*HI and *Pme*I and cloned into the *Bam*HI/*Pme*I sites of pEF6/myc-His (ThermoFisher Scientific). The plasmid obtained was named pEF-POLH. *POLH^−/−^* cells (1 × 10^6^) were transfected with 10 μg of ScaI-linearized pEF-POLH using a NEPA21 Super Electroporator (Nepa Gene Co., Ltd.) according to the manufacturer’s instructions. After incubating for 48 h in RPMI-1640 medium, cells were plated into 96-microwell plates in the presence of 10 μg/ml blasticidin S. Blasticidin S-resistant clones expressing *POLH* were then separated. The *XPA-* and POLH-expressing clones were then subjected to UV irradiation and *supF* forward mutation assay.

### UV irradiation

Cells in the exponential phase were washed and resuspended with RPMI-1640 medium without phenol red prior to the irradiation. Moreover, cells (2.5 × 10^6^) suspended in 5 ml medium were placed in a 100 mm petri dish and were exposed to UVC light. After irradiation, cells were cultured for 3 days. The relative growth rate was determined by counting cells over the 3 day period using Coulter Counter Z2 (Beckman).

### *supF* forward mutation assay

The position 112 of the *supF* gene was selected as the site for embedding rG, 8-oxo-rG, or 8-oxo-dG according to the previous study^55^. The 5’-phosphorylated 21-mer primer (5’-CGACTTCGAAGNTTCGAATCC-3’, where N represents dG rG, 8-oxo-rG, or 8-oxo-dG) was synthesized and provided by Tsukuba Oligo Service Co., Ltd. (Ibaraki, Japan). Each primer was annealed with pMY189 single-stranded DNA that was prepared using the VCSM13 helper phage. The closed circular double-stranded DNA, containing a single dG, rG, 8-oxo-rG, or 8-oxo-dG, was then synthesized and purified with ultracentrifugation as previously described^56^.

This closed circular double-stranded DNA (500 ng) was introduced into 2 × 10^6^ cells using NEPA21 Super Electroporator (Nepa Gene Co., Ltd.) according to the manufacturer’s instructions. After incubating for 48 h, the replicated plasmid was isolated from the cells and digested with *DpnI* to remove the unreplicated plasmids^57^. The resulting plasmid was introduced into the *E. coli* KS40/pOF105 indicator strain. To select *supF* mutants, the transformed cells were plated on LB plates containing nalidixic acid (Nal, 50 μg/ml), streptomycin (Sm, 100 μg/ml), ampicillin (Amp, 150 μg/ml), chloramphenicol (Cm, 30 μg/ml), X-Gal (80 μg/ml), and IPTG (23.8 μg/ml). For determining the total number of transformants, the transformed cells were plated onto LB plates containing Amp (150 μg/ml) and Cm (30 μg/ml). The frequency of *supF* mutants and mutation spectrum was determined as described previously^56^.

## Acknowledgements

We thank Dr. Tomonari Matsuda (Kyoto University) for providing us with pMY189 and Dr. Tatsuo Nunoshiba (International Christian University) for providing us with KS40/pOF105. We are grateful to Dr. Samuel H. Wilson (NIEHS, National Institutes of Health) and Dr. Masayuki Yokoi (Kobe University) for helpful comments and discussions. We wish to thank Dr. Takeshi Yasuda (National Institute of Radiological Sciences) for allowing us to use the UV irradiator. We thank Enago (www.enago.jp) for English-language review. This work was supported by the joint research program of the Biosignal Research Center, Kobe University [291002 to A.S.] and JSPS KAKENHI [19K12339 and 16K16195 to A.S., 25281022 to M.Y., and 16H02953 to H.S.]. This work was also supported by the grants from the Takeda Science Foundation [to K.U. and H.S.] and Mitsubishi research foundation [to H.S.].

## Author Contributions

A.S. and H.T. designed the research, A.S., H.T., A.T., K.H., M.S., M.T., H.S., S.T., K.S., M.Y., M.H., and K.U. discussed the study and set up the experiments. A.S., H.T., A.T., K.H., M.S., and M.T. performed the research and analyzed data. A.S. and H.T. wrote the paper. All authors read and approved the final manuscript.

