## Supplementary Figure S1 for "Processing of a single ribonucleotide embedded into DNA by human nucleotide excision repair and DNA polymerase η"

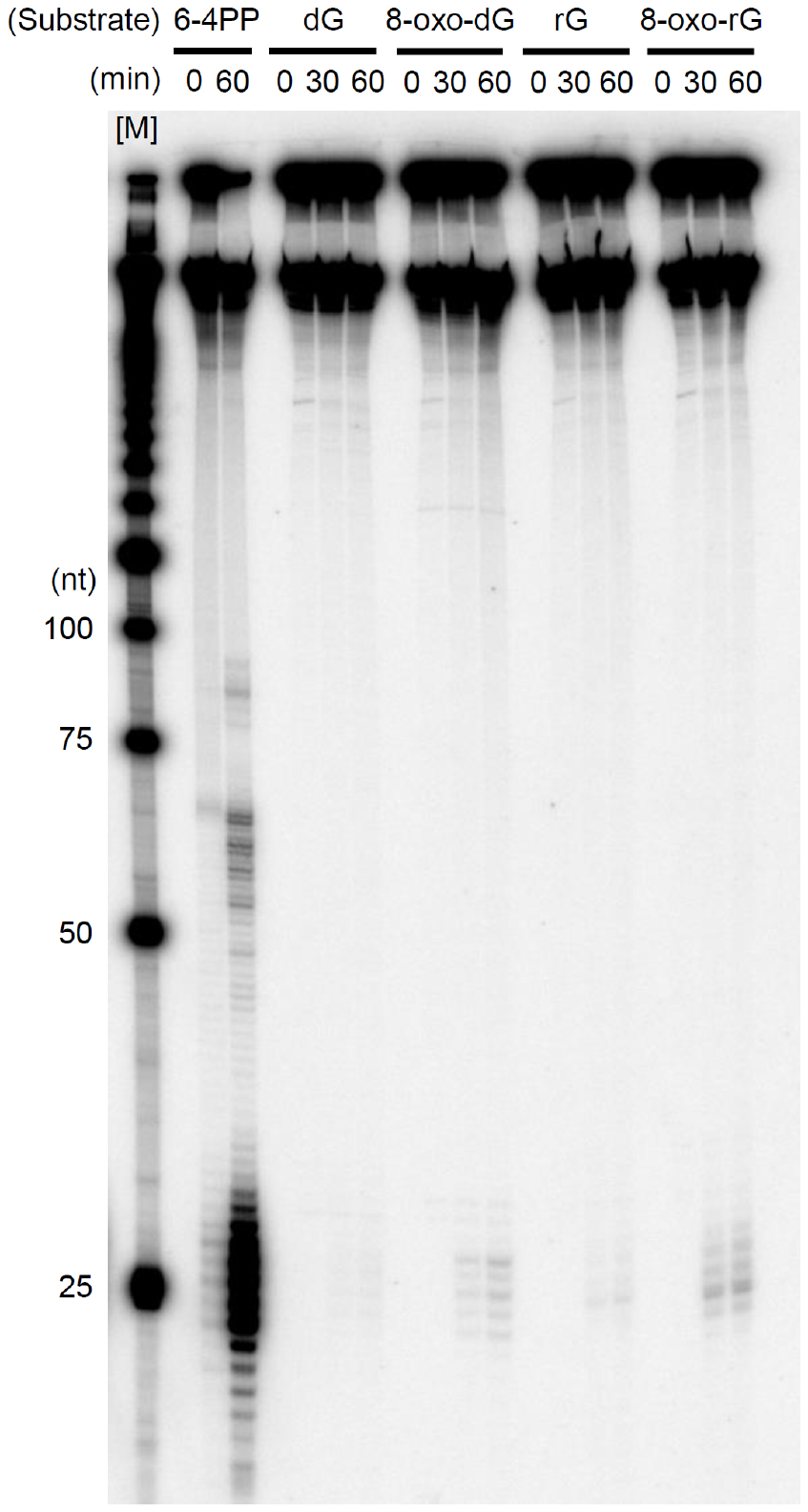

**Figure S1** Reconstituted human nucleotide excision repair excises oxidized ribonucleotide. The substrates containing UV-induced 6-4-photoproducts (6-4PP), dG, 8-oxo-dG, rG, and 8-oxo-rG were internally labeled with ^32^P and used for the dual incision assay. The DNA samples were subjected to 10% denaturing polyacrylamide gel electrophoresis followed by autoradiography. [M] ^32^P-labeled 25-nucleotides (nt) ladder.
